# *Escherichia coli* ZipA organizes FtsZ polymers into dynamic ring-like protofilament structures

**DOI:** 10.1101/319228

**Authors:** Marcin Krupka, Marta Sobrinos-Sanguino, Mercedes Jiménez, Germán Rivas, William Margolin

## Abstract

ZipA is an essential cell division protein in *Escherichia coli*. Together with FtsA, ZipA tethers dynamic polymers of FtsZ to the cytoplasmic membrane, and these polymers are required to guide synthesis of the cell division septum. This dynamic behavior of FtsZ has been reconstituted on planar lipid surfaces *in vitro*, visible as GTP-dependent chiral vortices several hundred nm in diameter, when anchored by FtsA or when fused to an artificial membrane binding domain. However, these dynamics largely vanish when ZipA is used to tether FtsZ polymers to lipids at high surface densities. This, along with some *in vitro* studies in solution, has led to the prevailing notion that ZipA reduces FtsZ dynamics by enhancing bundling of FtsZ filaments. Here, we show that this is not the case. When lower, more physiological levels of the soluble, cytoplasmic domain of ZipA (sZipA) were attached to lipids, FtsZ assembled into highly dynamic vortices similar to those assembled with FtsA or other membrane anchors. Notably, at either high or low surface densities, ZipA did not stimulate lateral interactions between FtsZ protofilaments. We also used *E. coli* mutants that are either deficient or proficient in FtsZ bundling to provide evidence that ZipA does not directly promote bundling of FtsZ filaments *in vivo*. Together, our results suggest that ZipA does not dampen FtsZ dynamics as previously thought, and instead may act as a passive membrane attachment for FtsZ filaments as they treadmill.

**IMPORTANCE:** Bacterial cells use a membrane-attached ring of proteins to mark and guide formation of a division septum at mid-cell that forms a wall separating the two daughter cells and allows cells to divide. The key protein in this ring is FtsZ, a homolog of tubulin that forms dynamic polymers. Here, we use electron microscopy and confocal fluorescence imaging to show that one of the proteins required to attach FtsZ polymers to the membrane during *E. coli* cell division, ZipA, can promote dynamic swirls of FtsZ on a lipid surface *in vitro*. Importantly, these swirls are only observed when ZipA is present at low, physiologically relevant surface densities. Although ZipA has been thought to enhance bundling of FtsZ polymers, we find little evidence for bundling *in vitro*. In addition, we present several lines of *in vivo* evidence indicating that ZipA does not act to directly bundle FtsZ polymers.

## INTRODUCTION

Bacterial septation is a complex process and dozens of essential and accessory proteins participate to assemble the cell division machinery, the divisome. In *Escherichia coli* the earliest event in the septum formation is the assembly of FtsZ, FtsA and ZipA into the proto-ring, a discontinuous structure at mid-cell that serves as a scaffold for the rest of the divisome components (1, 2).

FtsZ, a prokaryotic tubulin homologue, assembles into GTP-dependent protofilaments required for divisome activity (3-7). These FtsZ filaments are anchored to the inner surface of the cytoplasmic membrane by both FtsA and ZipA, and migrate in patches around the cell circumference by treadmilling. Through connections involving other divisome proteins that cross the cytoplasmic membrane, these treadmilling FtsZ protofilaments help to guide the septum synthesis machinery in concentric circles, resulting in inward growth of the septal wall until it closes and the daughter cells are separated (8, 9).

Although FtsA is conserved throughout diverse bacterial species, ZipA is limited to gamma-proteobacteria, including *E. coli* (10). In the absence of both FtsA and ZipA, FtsZ fails to attach to the membrane or form the proto-ring, demonstrating the requirement for a membrane tether (11). In the presence of only FtsA or ZipA, FtsZ filaments form a membrane-anchored ring, but septation fails to proceed (12), suggesting that the divisome is in a locked state. One major unanswered question in the field is why *E. coli* requires dual FtsZ membrane anchors to assemble a divisome that completes septation. Our recent study provides a potential answer by showing that FtsA exerts a specific structural and functional constraint on FtsZ protofilaments: when attached to lipid monolayers, FtsA assembles into clusters of polymeric minirings that align FtsZ polymers and inhibit their bundling (13).

In this report we use the term “bundling” in reference to increased lateral interactions between adjacent FtsZ protofilaments, resulting in two or more polymers closely associated in parallel. The physiological role of these lateral interactions is not firmly established, but several FtsZ mutants that are defective in protofilament bundling *in vitro* are also defective in cell division (14-16). In addition to the intrinsic ability of FtsZ polymers to interact laterally, proteins called Zaps (ZapA, C, D; FtsZ-associated proteins) help to bundle or crosslink FtsZ polymers *in vitro* (17, 18). Inactivation of single Zap proteins is not lethal, but mutant cells lacking multiple Zap proteins have significant division defects (19-23). Hyper-bundled mutants of FtsZ have also been isolated, and cells expressing these alleles also divide abnormally (24-26). However, one hyper-bundling mutant, called FtsZ*, has gain-of-function properties (27). FtsZ*, which forms mostly double stranded filaments *in vitro*, allows division of cells lacking ZipA and can resist the effects of other FtsZ inhibitors. Together, these findings suggest that lateral interactions are important for FtsZ function, but these interactions need to be balanced.

The aforementioned study (13) proposed a model in which FtsA minirings antagonize FtsZ protofilament bundling, keeping the divisome in a locked state. In this model, once the cell is ready to divide, these minirings are disrupted and are no longer a constraint for FtsZ polymer bundling. This is consistent with another model in which broken FtsA polymers start to recruit later divisome components, while FtsZ polymers become anchored to cell membrane by ZipA (2, 28). ZipA has been shown to stabilize the proto-ring, not only by anchoring FtsZ to the membrane, but also by protecting it from degradation by ClpXP protease (29-31). Whereas FtsA inhibits FtsZ polymer bundling (13), ZipA is considered an FtsA competitor for FtsZ polymers because of their common binding site at the FtsZ C terminus (32-35). Thus, it is not surprising that ZipA has been suggested as a bundler of FtsZ. However, the reports on its effect on FtsZ protofilament bundling in solution are not consistent (27, 36-40).

Recently it has become clear that the functionalities of the proto-ring proteins need to be tested in a more physiological context by attaching them to a lipid surface (13, 41-47). For example, Mateos-Gil et al. (39) used atomic force microscopy to visualize FtsZ polymers bound to *E. coli* lipid bilayers through ZipA. These ZipA-tethered FtsZ molecules formed a dynamic two-dimensional network of curved, interconnected protofilaments that seemed to be bundled. On the contrary, ZipA incorporated into phospholipid bilayer nanodiscs did not trigger significant FtsZ polymer bundling (29). Finally, Loose and Mitchison (44) reconstituted the *E. coli* proto-ring components on supported lipid bilayers and showed that FtsA organized FtsZ polymers into dynamic patterns of coordinated streams and swirling rings with preferential directions, which suggested treadmilling. Importantly, these dynamics were sharply reduced when FtsZ protofilaments were attached to the membrane by ZipA or when using artificially membrane-targeted FtsZ. Although the resulting FtsZ polymers were described as bundled, the resolution obtained by TIRFM probably could not distinguish between single and bundled FtsZ protofilaments. More recently, it was found that artificially membrane-bound FtsZ self-organizes into similar vortices, even in the absence of FtsA (45). This effect casts doubt on the dampening effects of ZipA on FtsZ dynamics observed previously.

In this study, we revisit the effect of ZipA on FtsZ protofilaments, including its role in polymer bundling. In contrast to the prevailing model, our *in vivo* results show that unlike Zaps, FtsZ* or the FtsA* gain of function mutant (48), ZipA does not play a significant role in FtsZ protofilament bundling. We further show that as previously reported (46, 49) (Sobrinos-Sanguino et al., in preparation), the surface concentration of ZipA is critical in controlling the activities and interactions with FtsZ *in vitro*. Using a His_6_-tagged soluble variant of ZipA (sZipA) immobilized on lipids, we demonstrate that this protein organizes FtsZ into similar swirling vortices of mostly single protofilaments, a role that was previously attributed exclusively to FtsA (44). These results provide further evidence that ZipA does not inhibit FtsZ polymer dynamics at the membrane.

## RESULTS

### FtsA* or excess FtsZ rescue the FtsZ bundling deficient ∆*zapA*∆*zapC* mutant

Recently we reported that FtsZ protofilament bundling is antagonized by FtsA, both *in vivo* and *in vitro* (13). For example, FtsA overproduction reverses the toxic effects of FtsZ over-bundling triggered by excess ZapA. Conversely, even a slight excess of FtsA exacerbates the already moderately filamentous cell phenotype of ∆*zapA* or the more severe filamentous phenotype of ∆*zapA*∆*zapC* double mutants (13), which lack one or two FtsZ bundling proteins, respectively (18). We also previously reported that the self-bundling mutant *ftsZ\** (encoding (FtsZ_L169R_) could completely suppress the cell division deficiency of the ∆*zapA*∆*zapC* double mutant (27). This suggested that if FtsZ protofilaments are bundled by factors independent of ZapA and ZapC, then the requirement for the latter proteins in normal cell division could be bypassed.

We first surmised that a moderate increase in intracellular FtsZ concentration could promote polymer bunding simply by molecular crowding, and this could bypass the need for ZapA and ZapC and consequently suppress the filamentation of many ∆*zapA*∆*zapC* cells (Fig. 1A-B). To produce extra FtsZ, we used the pJF119HE-FtsZ plasmid (26). As expected, FtsZ overproduction suppressed ∆*zapA*∆*zapC* cell filamentation (Fig. 1C), supporting the idea that higher FtsZ protein concentration favors increased lateral interactions between protofilaments.

**Figure 1.**
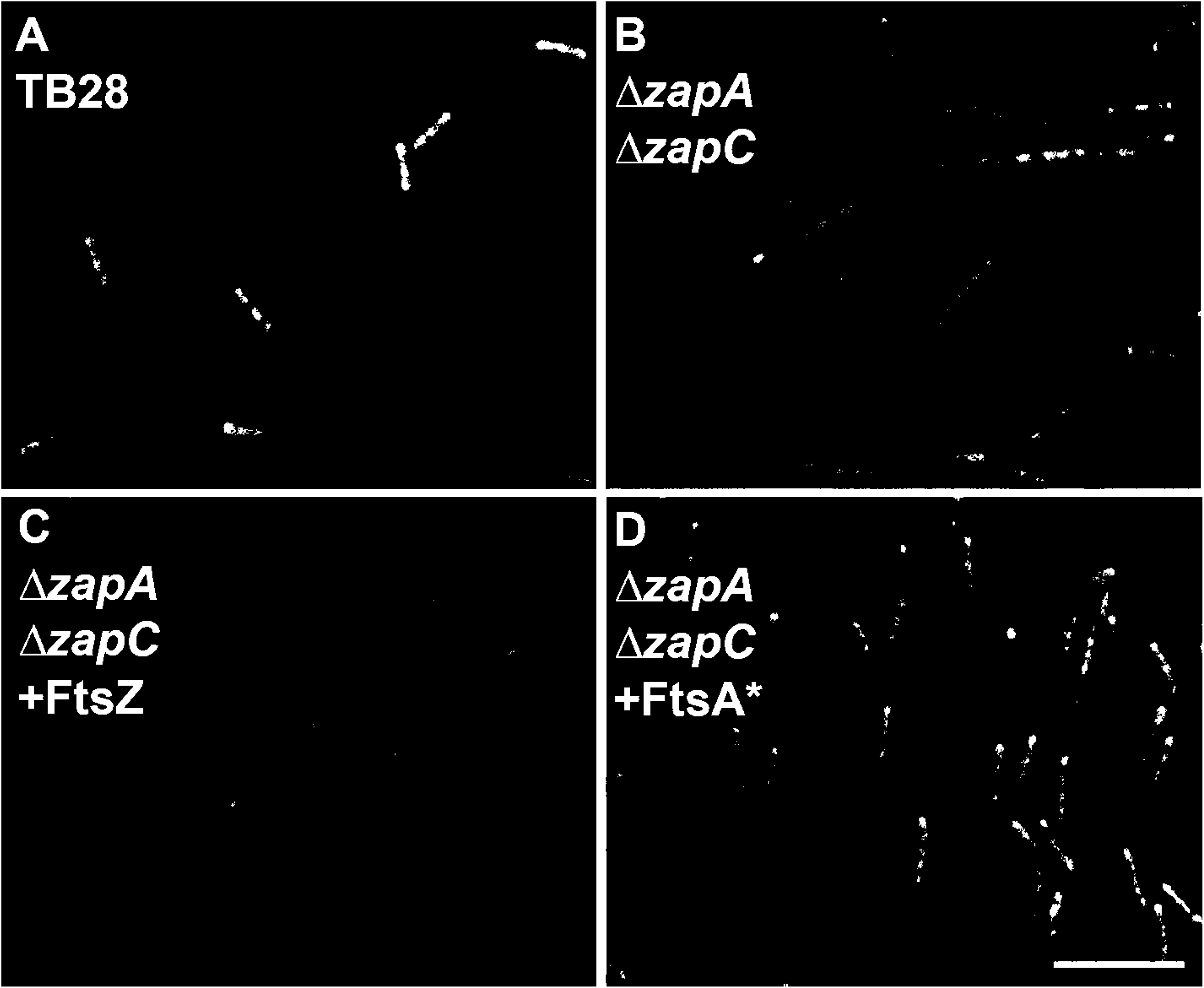
FtsA* or excess FtsZ can rescue the FtsZ bundling deficient ∆*zapA*∆*zapC* mutant. (A) Cells of the TB28 parent, or (B-D) ∆*zapA*∆*zapC* double mutant carrying either no plasmid (B), pJF119HE-FtsZ (C) or pDSW210F-FtsA* (D) were grown to mid-logarithmic phase in LB medium (supplemented with 50 μM IPTG in C) and imaged using DIC. Scale bar= 10 μm.

We then asked whether FtsA*, a potent gain-of-function mutant that repairs multiple cell division defects (48, 50-53), could bypass the need for ZapA and ZapC. Unlike wild-type FtsA, FtsA* promotes FtsZ polymer bundling on lipid monolayers (13). We found that even uninduced levels of FtsA* from pDSW210F-FtsA* were sufficient to completely rescue the division defects of ∆*zapA*∆*zapC* cells (Fig. 1D). Therefore, FtsA* has the same rescuing effect as FtsZ* in the absence of ZapA and ZapC, supporting the idea that FtsA*, ZapA and ZapC all promote FtsZ protofilament bundling like FtsZ*.

### Excess ZipA cannot counteract cell division defects caused by deficient FtsZ bundling

As already mentioned, *E. coli* FtsA inhibits FtsZ polymer bundling. Our *in vitro* results indicate that this occurs due to the unusual mini-ring polymers that purified FtsA forms on lipid monolayers. In contrast, purified FtsA* forms shorter curved oligomers under similar conditions. Although the mechanism is not yet known, these FtsA* arcs no longer inhibit FtsZ polymer bundling and instead permit or promote it, consistent with our *in vivo* results (Fig. 1D). This was most apparent when FtsA* was combined with FtsZ* on lipid monolayers: in a striking additive effect, large sheets were formed consisting of many laterally associated protofilaments (13). Interestingly, both gain of function mutants that promote FtsZ bundling, FtsA* and FtsZ*, bypass the need of the third proto-ring component, ZipA (27, 48). This, along with evidence that purified ZipA can bundle FtsZ under certain conditions, led to the hypothesis that ZipA might also trigger FtsZ bundling (54).

In this scenario, excess ZipA should be able to rescue the ∆*zapA*∆*zapC* cell filamentation phenotype, similarly to excess FtsZ, FtsA* or FtsZ* (27) (Fig. 1). To test this, we first transformed the ∆*zapA*∆*zapC* double deletion and its TB28 wild-type parental strain (55) with pKG110-ZipA, a plasmid that expresses *zipA* from a salicylate-inducible *nahG* promoter and a weak ribosome binding site that keeps expression low. Notably, uninduced levels of ZipA from pKG110-ZipA did not suppress the ∆*zapA*∆*zapC* filamentous phenotype (Fig. 2B). Instead, the induction of *zipA* expression was consistently more toxic not only for ∆*zapAzapC*, but also for the ∆*zapA* single deletion strain when compared with the wild-type parent TB28 (Fig. 2A). Endogenous FtsZ was produced at similar levels in both ZipA-uninduced and induced cells, ruling out the possibility that excess ZipA could affect viability through changes in FtsZ intracellular levels (Fig. S1A).

**Figure 2.**
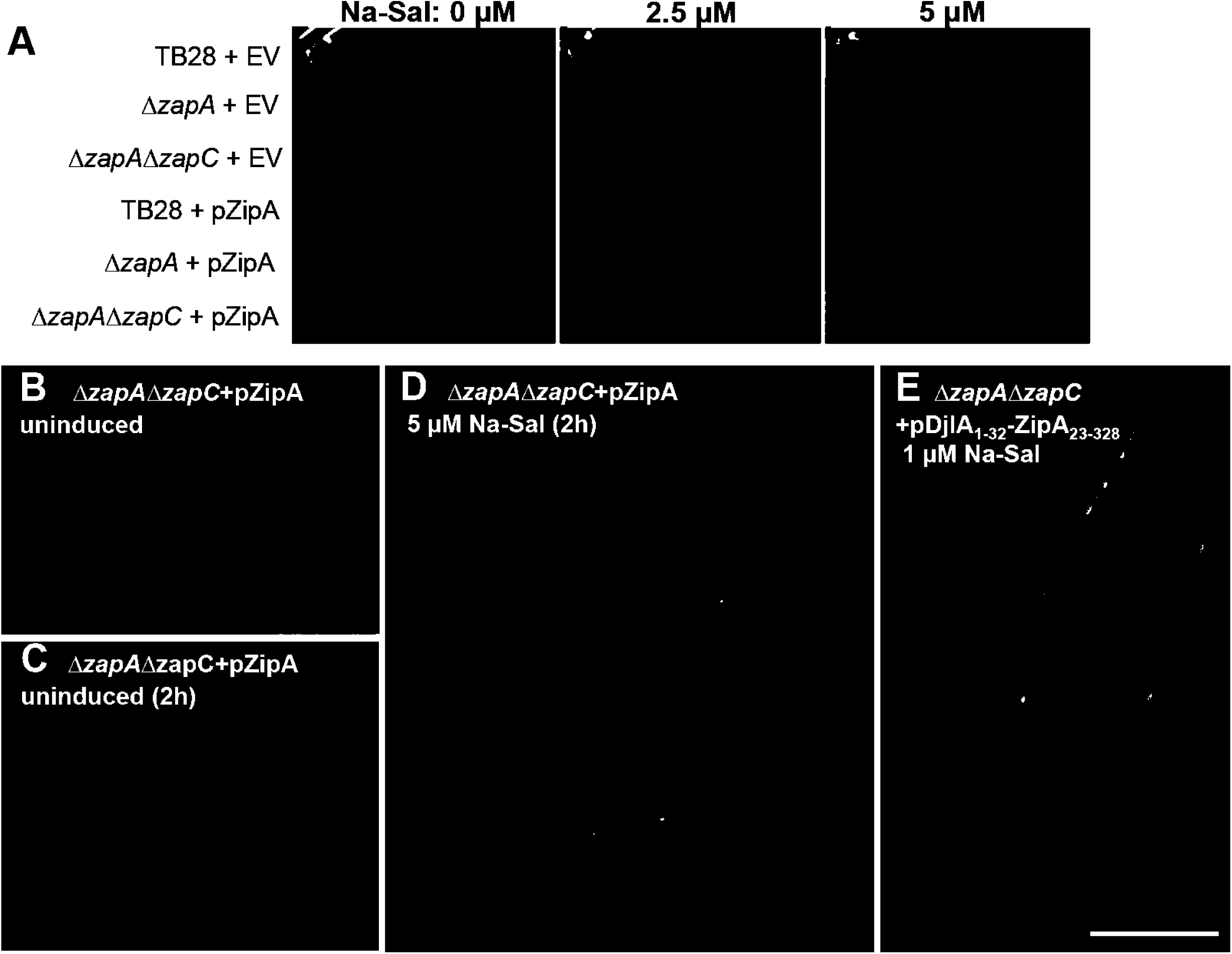
Excess ZipA or a ZipA with a swapped transmembrane domain cannot compensate for cell division deficiencies of ∆*zap* mutants. (A) TB28 and Zap-deletion strains ∆*zapA* and ∆*zapA*∆*zapC* transformed with pKG110 empty vector (EV) or pKG110-ZipA (pZipA) were grown to exponential phase and plate spotted IN 10-fold dilutions at different concentrations of inducer (sodium salicylate; Na-Sal). (B-E) Representative DIC images of ∆*zapA*∆*zapC* cells transformed with pKG110-ZipA (B-D) and pKG116-DjlA_1___32_-ZipA_2_3–_328_ (E) are shown (see panel descriptions and text). Scale bar= 10 ^m.

Further growth until late exponential phase exacerbated the already elongated cell phenotype of the ∆*zapA*∆*zapC* strain both in the absence (Fig. 2C) and presence of inducer (Fig. 2D); cells of the ∆*zapA* single mutant behaved similarly (not shown). These results suggest that ZipA might not be a bundler of FtsZ polymers, contrary to what we initially expected.

The region of ZipA known to interact with FtsZ polymers is the FZB (FtsZ Binding) globular domain at its C-terminal end (34, 37, 56, 57). To exclude the possibility that the toxicity of excess ZipA for ∆*zapA*∆*zapC* cells was due to the accumulation of transmembrane domains at septation sites (49) or because the N-terminal transmembrane region of ZipA might affect cell division by an unknown mechanism, we separated the FZB domain from the transmembrane region. For this purpose we used a chimeric construct containing the C-terminal part of ZipA lacking the transmembrane region (ZipA_23-328_) fused to the N-terminal transmembrane domain of DjlA, (DjlA_1-32_), a protein not related to cell division (37). This hybrid membrane protein containing FZB was cloned into pKG116, a plasmid similar to pKG110 but with a stronger ribosome binding site for increased gene expression. However, similarly to the intact ZipA protein, the DjlA_1-32_-ZipA_23-328_ (FZB) protein was toxic and exacerbated the phenotype of ∆*zapA*∆*zapC* (Fig. 2E) and ∆*zapA* (not shown) cells. This further suggests that binding of ZipA to FtsZ polymers does not promote their bundling, or at least the type of bundling that could compensate for the lack of ZapA and ZapC (18).

To test the model further, we asked whether excess ZipA could rescue the dominant negative effects of an FtsZ allele (FtsZ_R174D_) that was reported to be defective in polymer bundling (14). Although a subsequent study suggested that FtsZR174D was capable of bundling under certain conditions (58), we recently confirmed (Schoenemann et al., in revision) that this protein is indeed more bundling-defective than wild-type FtsZ, as suggested in the original report.

To test this idea, we constructed a strain with two plasmids: pDSW210F-ZipA-GFP, and either pKG110-FtsZ or pKG110-FtsZ_R174D_, such that expression of ZipA-GFP is controlled by IPTG and expression of the FtsZ derivatives is controlled by sodium salicylate. The ZipA-GFP is functional and can complement a *zipA1*(ts) mutant (59). Expression of FtsZ_R174D_ at any level above 1 μM sodium salicylate was strongly dominant negative (Fig. S2), in contrast to FtsZ, which allowed viability even at 2.5 μM (and higher, not shown). Notably, ZipA, whether uninduced or induced with IPTG, was unable to counteract the dominant negative effects of FtsZ_R174D_, consistent with the idea that ZipA does not promote FtsZ bundling (Fig. S2). This is in sharp contrast with hyper-bundled FtsZ*, which is able to suppress the dominant negative effects of FtsZ_R174D_ (Schoenemann et al., in revision). Interestingly, the toxicity of ZipA at IPTG concentrations above 50 μM was antagonized by extra FtsZ. One possible explanation is that increased FtsZ bundling triggered by its higher intracellular levels (Fig. 1C) counteracts the negative effects of excess ZipA (Fig. S2).

### FtsA* and FtsZ* confer at least 10-fold resistance to excess ZipA

In our recent studies, we demonstrated that FtsZ* has an intrinsic capacity to bundle compared with wild-type FtsZ (27), whereas FtsA* can promote bundling of wild-type FtsZ protofilaments (13). Moreover, both gain of function mutants correct the defective division phenotype of ∆*zapA*∆*zapC* under-bundling mutants (Fig. 1). If ZipA acts to bundle FtsZ polymers, its excess in an *ftsZ\** or *ftsA\** background should result in over-bundling and be toxic for the cells by inhibiting cell division, as previously reported (Haeusser et al., 2015). To test this idea, we transformed WM1659 and WM4915, which replace the native chromosomal *ftsA* or *ftsZ* with *ftsA\** or *ftsZ\** alleles, respectively, with pKG110-ZipA in the WM1074 (MG1655) strain background. We found that ZipA overproduction from the *nahG* promoter was toxic at 5 μΜ sodium salicylate in the wild-type parent strain, and became more toxic at 10 μΜ inducer (Fig. 3, row 1). In contrast, the presence of *ftsA\** in WM1659 conferred full resistance against excess ZipA (Fig. 3, row 3), consistent with the original report (48). The effects of *ftsZ\** in WM4915 were more modest, but nonetheless resulted in at least a 10-fold increase in resistance at 5 μΜ inducer (Fig. 3, row 2). The effects of *ftsA*, ftsZ*\* or ZipA levels on viability were not due to changes in FtsZ levels, as these remained unchanged in the various conditions (Fig. S1B). The ability of alleles that promote FtsZ protofilament bundling to antagonize ZipA toxicity instead of exacerbate it is yet another argument against the idea that ZipA is a bundler of FtsZ.

**Figure 3.**
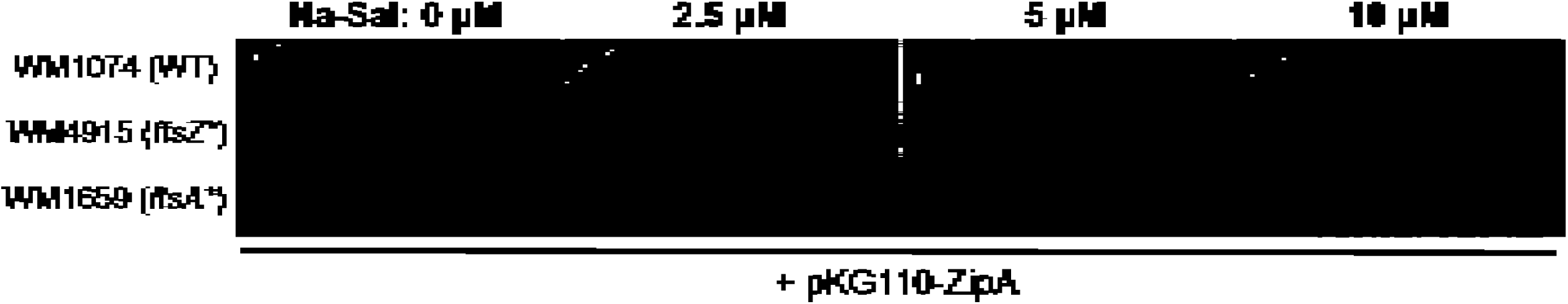
The *ftsZ\** or *ftsA\** alleles counteract the toxicity of excess ZipA. WM1074 wild-type strain and its derivatives containing chromosomal *ftsA\** (WM1659) and *ftsZ\** (WM4915), transformed with pKG110-ZipA were grown to exponential phase and spotted on plates at 10-fold dilutions with different concentrations of inducer (sodium salicylate; Na-Sal).

### Excess ZapA and ZapC only partially suppress the thermosensitivity of *zipA1*(ts)

We further explored whether ZipA has any functional overlap with Zap proteins by testing if their overproduction could rescue a thermosensitive *zipA1*(*ts*) mutant (12). We introduced plasmids expressing *ftsZ\** (positive control), *zapA, zapC, zapD*, or a combination of *zapA + zapC, zapA + zapD*, or *zapC + zapD* genes into the *zipA1*(*ts*) strain WM5337. As expected, FtsZ*, ZapA, ZapC, or ZapD all became toxic when overproduced (Fig. 4A), and only *ftsZ\** could fully suppress *zipA1*(*ts*) at 42°C (27) (Fig. 4C). This suggests that the FtsZ bundling promoted by Zaps cannot substitute for the absence of functional ZipA.

**Figure 4.**
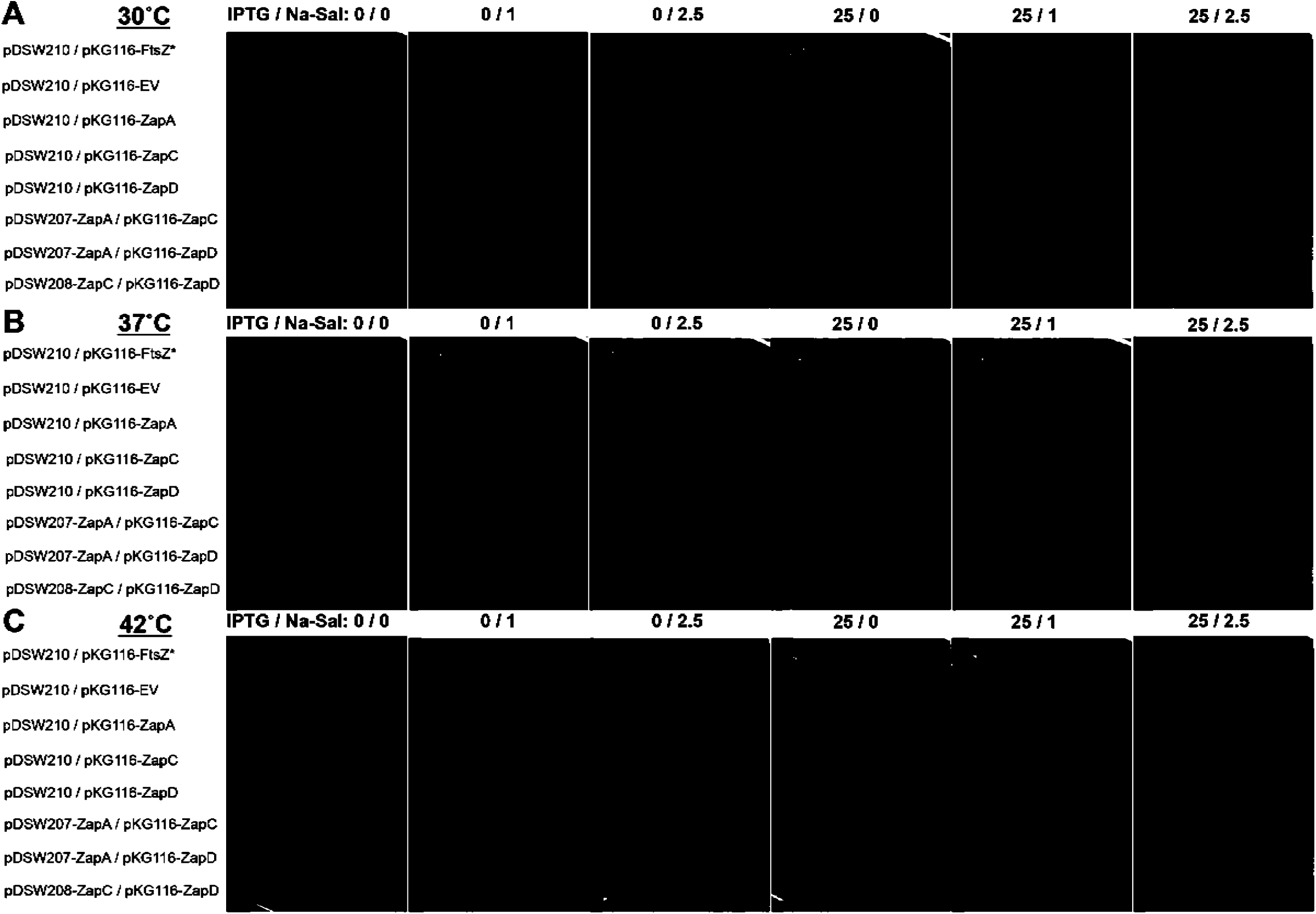
Excess Zap proteins only weakly suppress the *zipA1* thermosensitive allele. The WM5337 *zipA1* thermosensitive strain was transformed with the pairs of compatible plasmids indicated at each row, and spotted on pre-warmed plates at 10-fold dilutions containing indicated concentrations of inducers (IPTG for pDSW plasmids and Na-Sal for pKG116) and incubated at 30, 37 and 42°C.

The *zipA1*(*ts*) strain is also inviable at 37°C, and some factors can suppress the thermosensitivity of *zipA1* at these lower temperatures, including inactivation of certain amino acid biosynthesis genes (59). This suggests that the ZipA1 protein is partially active at 37°C, although not sufficient to sustain viability. To give the Zap proteins the best chance of suppressing *zipA1*, we tested whether the Zap proteins might be able to partially compensate for a partially defective ZipA at this less stringent temperature. We found that neither ZapD nor ZapA were able to suppress *zipA1* thermosensitivity at 37°C, but ZapC was (Fig. 4B). We also noticed a weak synergistic effect upon coexpression of both ZapA and ZapC, where there was a limited level of viability even at 42°C (Fig. 4C). Moreover, *zapA + zapC, zapA + zapD*, or *zapC + zapD* pairs also conferred partial suppression of *zipA1* thermosensitivity at 37°C (Fig. 4B). These results indicate that the Zap proteins and ZipA may have weak overlapping roles in FtsZ protofilament bundling, perhaps by enhancing the stability of the proto-ring and its tethering to the membrane and to the nucleoid (60).

### Low surface density ZipA organizes FtsZ into circular protofilament structures on lipid monolayers

So far, our *in vivo* data presented here are not consistent with the previous data that suggested ZipA is a major enhancer of FtsZ protofilament bundling. This prompted us to test whether ZipA had any effect on FtsZ bundling in an *in vitro* membrane system. For this, we examined the properties of FtsZ polymers on lipid monolayers. To date, this assay has been mainly used to visualize oligomeric structures of FtsZ protofilaments along with their FtsA membrane tethers by electron microscopy (13, 35, 43, 61, 62). Whereas FtsA has a short C-terminal amphipathic helix that acts as a membrane anchor (11, 61, 63), ZipA has a short N-terminal periplasmic region followed by a transmembrane domain (36, 37, 64).

Consequently, full-length ZipA could not be used in our assay.

Therefore, we decided to use an N-terminally truncated ZipA (soluble ZipA, sZipA) replacing the first 25 amino acids with an N-terminal His_6_ tag (65). To attach sZipA to the lipid monolayer, input lipids were supplemented with a nickel-chelating lipid (DGS-NTA) that anchors the His6 tag, thus mimicking the membrane topology of the full-length protein (44, 49, 65). The density of sZipA on the lipid monolayer surface was tuned by controlling the amount of NTA lipids added, as these two values are linearly proportional (Sobrinos-Sanguino, Ritcher and Rivas, in preparation). Importantly, we lowered the surface density of ZipA compared with previous studies (39, 44) by using 0.5-1% of NTA lipids instead of 10%, which more closely mimic physiologically relevant levels of ZipA. 0.5% NTA corresponds to a surface density of ~2000 ZipA molecules per μm^2^. Unperturbed *E. coli* cells contain ~1500 ZipA molecules per cell (66), which corresponds to around 400 molecules per μm^2^ assuming a uniform distribution. If 30% of these ZipA molecules are in a midcell ring that comprises 5-10% of the cell length, the estimated protein concentration in the ring would be ~2000 molecules per μm^2^, which is the low surface density we used.

As expected, when FtsZ was added without sZipA and examined by negative stain transmission EM, FtsZ polymers were scattered sparsely on the lipid monolayer, consistent with the requirement for a membrane anchor such as FtsA or ZipA (1). This residual binding was likely a result of random association of the FtsZ from the added solution onto the grid (Fig. S3B). However, when FtsZ was added to monolayers coated with low-density sZipA, we observed extensive FtsZ protofilament patterns. Most notably, these patterns differed depending on the concentration of NTA lipids, which in turn dictated the concentration of ZipA on the monolayer. For example, when FtsZ (1-5 μM) was polymerized with non-hydrolyzable GTP (GMPcPP) on monolayers seeded with low-density ZipA (0.5% NTA lipids out of the total input lipids), it became strikingly organized into circular structures of mostly single protofilaments in a repetitive pattern (Fig. 5A). These circular structures had an average of nine filaments per polymer. The external diameter was 279 ± 50 nm, with a lumen, lacking filaments, of ~100 nm in diameter. The lateral separation between the filaments was 10 ± 4 nm. The filaments that were closest together mostly appeared as double filaments, but were very loose and non-continuous (more than 70 structures were measured).

**Figure 5.**
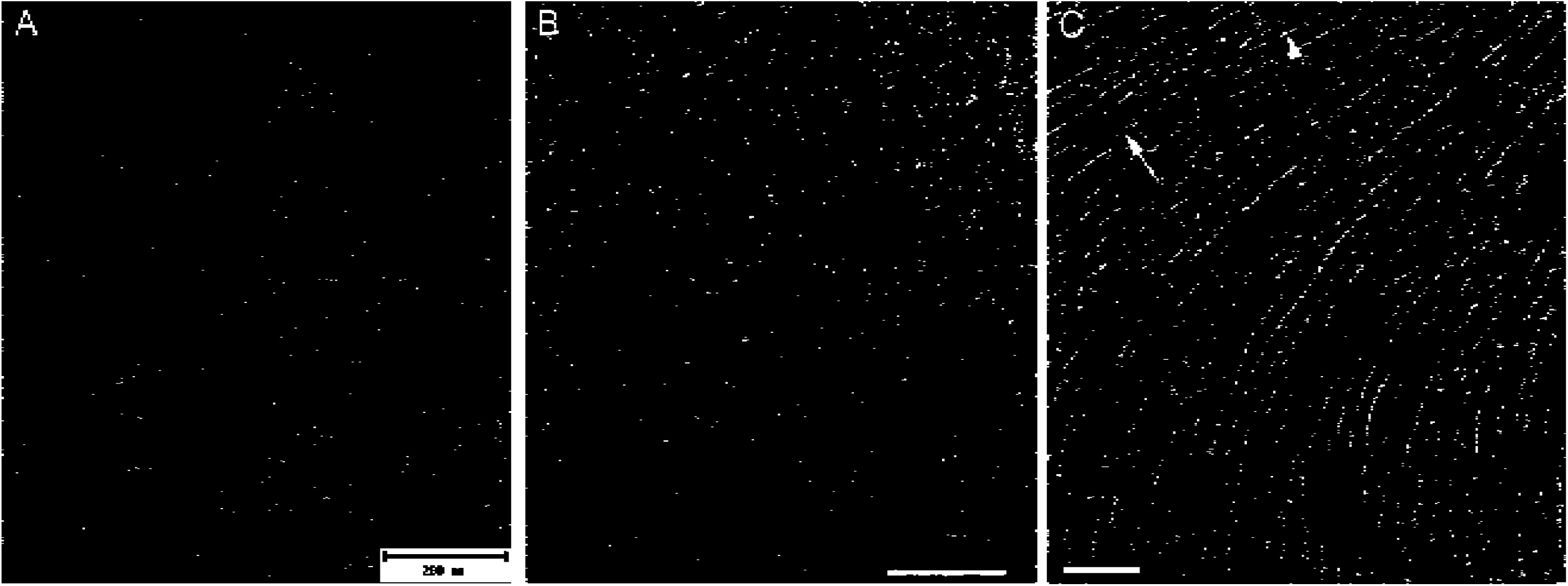
Assembly of FtsZ on *E. coli* polar lipid monolayers containing sZipA visualized by negative stain transmission electron microscopy. (A) Examples of circular structures of FtsZ single filaments on a lipid monolayer containing 0.5% DGS-NTA and sZipA (1 μΜ) in the presence of GMPcPP. FtsZ concentration was 1.5 μΜ. Scale bar = 200 nm. (B) Examples of circular structures of FtsZ single filaments on a sZipA-containing monolayer with 1% of DGS-NTA in the presence of GTP. FtsZ concentration was 2.5 μΜ. Scale bar = 200 nm. (C) Straight FtsZ filaments assembled in the presence of GTP on a lipid monolayer with a high surface density of sZipA (attached to 10% of DGS-NTA). The arrowhead highlights a typical single protofilament; the full arrow highlights a less common double protofilament. FtsZ and sZipA concentrations were 5 and 2 μΜ, respectively. Grids for all three panels were negatively stained and visualized by electron microscopy. Scale bar = 100 nm.

In the presence of GTP, which should support GTPase activity and filament treadmilling, the ring-like structures contained a smaller number of filaments (6 ± 2) but were larger than the structures formed in GMPcPP, with an external diameter of 400 ± 80 nm and a lumen 190 ± 20 nm in diameter. The GTP-FtsZ filaments appeared more separated than those formed with GMPcPP, as the average separation was 20 ± 9 nm (more than 50 structures were measured) (Fig. 5B). For both GTP and GMPcPP ring-like structures, the spacing measurements were compatible with the FtsZ filament arrangement found in the presence of FtsA minirings (13). To assess the effect of different lipids on these structures, we made lipid monolayers with DOPC. Similar ring-like structures containing FtsZ were observed with GTP (Fig. S4).

Next, we asked whether increasing surface density of sZipA might affect the ring-like structures of FtsZ polymers. We saw no difference between monolayers containing 0.5% vs. 1% NTA lipids (not shown). We then significantly increased the surface concentration of sZipA on monolayers by increasing the NTA concentration to 10%, mimicking ZipA overproduction *in vivo*. Whereas no oligomeric structures were detectable with sZipA alone (Fig. S3A), when FtsZ was added to the sZipA at this high surface density, polymers were strikingly aligned into parallel tracks of long, straight protofilaments spaced ~20 nm apart, and the formation of ring-like swirls observed at lower ZipA densities was abolished. Even at this high density of ZipA, most of FtsZ protofilaments remained unbundled (Fig. 5C). Whether FtsZ formed straight alignments or swirls was independent of FtsZ concentrations added to the reactions within the 1.5-5 μM physiological range (Fig. 5 and data not shown) (66).

### FtsZ swirls formed at low ZipA densities are highly dynamic and driven by GTP hydrolysis

The ring-like structures formed by FtsZ polymers on low density sZipA resembled the dynamic vortices formed either by FtsZ bound to membrane-attached FtsA or FtsZ fused to YFP and a membrane targeting sequence (FtsZ-YFP-mts) on supported lipid bilayers (44, 45). This prompted us to analyze the dynamics of fluorescently labeled FtsZ protofilaments on bilayers containing 0.5% NTA (low density sZipA) using confocal microscopy. When GTP was added to trigger FtsZ polymerization, we observed swirling vortices with a chiral clockwise rotation, similar to those from the aforementioned reports (Fig. 6A, Video S1). A consistently negative slope of the kymographs (Fig. S5) confirmed the directionality of the rotation within the rings. The estimated rotational speed within these structures was ~ 1.8 μm min^-1^. Similar, but markedly less dynamic swirling rings were observed in the presence of GMPcPP (Fig. 6B, Video S2). The estimated speed was 0.3 μm min^-1^, consistent with the idea that vortex formation is driven by GTP hydrolysis (45).

**Figure 6.**
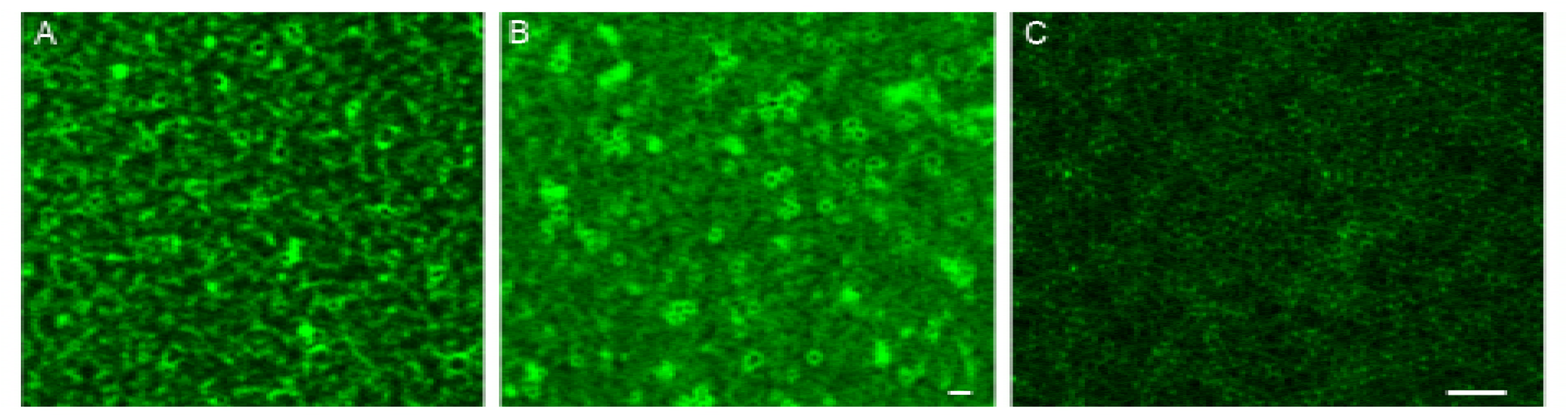
Fluorescence microscopic images of FtsZ assembly on SLBs of *E. coli* polar lipids containing sZipA. (A) Assembly of Alexa 488-labeled FtsZ into dynamic vortices after GTP addition. (B) Assembly of similar circular structures after GMPcPP addition. (C) Representative high resolution STED image of the experiment shown in panel B. FtsZ and sZipA concentrations were 1 μΜ. Scale bars = 1 μm.

To visualize the structure of these vortices in more detail, we used super-resolution microscopy (STED). These structures were sharper than those imaged by standard confocal microscopy and their size was similar to the size of the lipid monolayer-attached swirls observed previously by electron microscopy (Fig. 6C). We also used total internal reflection fluorescence microscopy (TIRFM) to visualize the FtsZ swirls at low ZipA surface density. We confirmed that these swirls formed both in GTP and GMPcPP (data not shown).

However, the TIRFM approach, which is highly sensitive to the distance of the fluorophore from the surface, was hampered by significant image fluctuation, most probably due to the movement of the unstructured domain of ZipA. This precluded a more precise analysis of the FtsZ swirls by TIRFM.

## DISCUSSION

Here, we provide *in vivo* and *in vitro* evidence that ZipA does not inhibit FtsZ treadmilling dynamics, unlike what was suggested previously (44). Instead, when FtsZ protofilaments are tethered to lipids by sZipA at levels that probably more closely mimic physiological conditions, they align and curve to form dynamic swirls that are very similar to those observed previously by FtsA-mediated tethering to lipids (44), direct adsorption to a mica surface (67, 68), or when subjected to crowding agents (69). These swirls, whose dynamics depend on GTP hydrolysis, likely represent treadmilling FtsZ polymers that comprise the FtsZ ring *in vivo* (8, 9). When we tested sZipA at an artificially high density on the lipid surface by applying a high (10%) concentration of NTA lipids, FtsZ protofilaments aligned into large, straight, apparently static structures that are micrometers in length. This observation is consistent with a previous study using high surface concentrations of sZipA, which concluded that ZipA curtails FtsZ dynamics (44). Therefore, we propose that lower surface ZipA densities are necessary to allow FtsZ protofilaments the needed flexibility for their characteristic dynamic movement along the membrane, which is crucial for guiding septum synthesis (8, 9).

Despite previous reports that ZipA bundles FtsZ when in solution, including stabilizing highly curved or circular forms of FtsZ polymers (40), here we present several lines of evidence that ZipA does not directly bundle FtsZ protofilaments at lower, probably more physiological densities on lipid surfaces or in *E. coli* cells. When attached to a lipid monolayer at these densities, sZipA efficiently tethers and aligns FtsZ protofilaments, but close lateral associations were uncommon. Even at high surface densities of sZipA that promoted extensive and relatively static FtsZ filament alignments, most protofilaments remain apart, indicating that ZipA does not directly bundle FtsZ like FtsA* does (13).

Furthermore, if ZipA actually stimulates FtsZ protofilament bundling, then it might be expected to replace the bundling functions of Zap proteins in cells. Instead, and in contrast to FtsA*, excess ZipA failed to rescue the cell division deficiency of ∆*zapA* or ∆*zapA*∆*zapC* mutant cells. ZipA also failed to counteract the dominant negative phenotype of the likely bundling-defective FtsZ_R174D_. In another test of ZipA’s bundling ability *in vivo*, it was predicted that excess ZipA might be more toxic in a bundling-proficient *ftsA\** or *ftsZ\** strain background compared with a normal background, due to FtsZ over-bundling. Instead, the *ftsA\** or *ftsZ\** alleles actually antagonized the toxicity of excess ZipA by at least 10-fold, suggesting again that ZipA is not acting significantly to bundle FtsZ. However, it is also possible that FtsA* and FtsZ* may have already maximally bundled the FtsZ in the cell, leaving no room for additional bundling by ZipA if it were to occur. The mechanism by which *ftsA\** or *ftsZ\** suppresses ZipA toxicity cannot yet be ascertained, as it is not yet known why excess ZipA is toxic.

These results suggest that ZipA is not a significant bundling factor for FtsZ, or at least that its mechanism of action is distinct from that of Zaps, FtsA* and FtsZ*. Nevertheless, extra ZapC could rescue the thermosensitivity of a *zipA1* mutant at 37°C, and even at the most stringent temperature of 42°C, a combination of ZapA and ZapC was able to rescue growth somewhat. One explanation for this is that crosslinking of FtsZ polymers by extra ZapA/ZapC generally promotes FtsZ protofilament alignment in parallel superstructures (i.e., swirls) that mimic the swirls assembled by ZipA, thus stabilizing the proto-ring. Because it is not clear what functions of the mutant ZipA1 protein are compromised at less stringent nonpermissive temperatures, it is difficult to know what ZapC is rescuing at 37°C that it cannot rescue at 42°C.

This brings up a broader question: why is ZipA essential for divisome function if it performs what seems to be a very similar function as FtsA? Both promote FtsZ protofilament alignment without permitting bundling *in vitro*, and their *in vivo* phenotypes are consistent with this, so why are both necessary *in vivo*? For example, when ZipA is inactivated, even in the presence of FtsA, recruitment of downstream divisome proteins is blocked, implicating ZipA in that essential function (70). We favor the idea that ZipA has additional roles in later divisome function that are distinct from those of FtsA. Furthermore, the ability of certain mutants such as FtsA* and FtsZ* to bypass ZipA may not be due solely to restoration of FtsZ bundling. For example, FtsA* likely recruits downstream divisome proteins more effectively than FtsA, and can accelerate cell division (28, 51, 71). It remains to be seen what these other activities of ZipA are and how they differ from the activities of FtsA. It was previously suggested that the ability of ZipA to form homodimers via its N-terminal domain might enhance FtsZ protofilament bundling (72). Although our lipid monolayer assays probably did not permit homodimerization of sZipA given that the native N terminus is missing, our genetic data using native ZipA suggest that its homodimerization does not significantly promote FtsZ bundling in vivo.

Another important question is how the FtsZ protofilaments become aligned as they self-assemble on lipids along with their membrane tethers. The study of plant microtubules may provide clues. During growth of the cortical microtubule array in plant cells, microtubules align with each other in a self-reinforcing mechanism. When a plus end of a microtubule meets another microtubule at an angle of less than 40°, the first polymer’s plus end changes direction and ends up parallel with the encountered polymer. When faced with another microtubule at angles greater than 40°, the plus end is more likely to disassemble (catastrophe), thus selecting against crossovers and reinforcing parallel alignments (73, 74). Such behavior, coupled with the tendency of intrinsically curved FtsZ protofilaments to adopt the intermediate curved conformation (67, 75, 76), could explain how the swirls become established and self-perpetuate. These curved groups of FtsZ polymers may be important to generate bending forces at the membrane (42, 75). It is possible that highly curved FtsZ also has a role in this activity, given that FtsZ minirings only ~25 nm in diameter can assemble on lipid monolayers (40). Although a specific type of membrane tether is not required for the generation of swirls (45), our data from this study and from our recent report (13) indicate that both FtsA and ZipA maintain FtsZ protofilaments in an aligned but mostly unbundled state. Yet the gain-of-function properties of FtsA* and FtsZ*, and their ability to specifically promote FtsZ lateral interactions, suggest that progression of the divisome requires a set of factors that ultimately switch FtsZ protofilaments to a bundled form. The ability of FtsA* and FtsZ* to bypass ZipA suggests that ZipA itself may be one of these factors, but that it does not necessarily act directly on FtsZ.

## MATERIALS AND METHODS

### Reagents

*E. coli* polar lipid extract (EcL), 1,2-dioleoyl-*sn*-glycero-3-phosphocholine (DOPC), and 1,2-dioleoyl-*sn*-glycero-3-[(N-(5-amino-1-carboxypentyl)iminodiacetic acid) succinyl] (nickel salt))(NTA) from Avanti Polar Lipids, Inc. (Alabaster, AL), were kept as 10-20 g/l stocks in chloroform solutions. Alexa Fluor 488 and Alexa Fluor 647 succinimidyl ester were from Molecular Probes/Invitrogen. GTP was from Sigma. GMPcPP (Guanosine-5'-[(α,β)-methyleno]triphosphate, Sodium salt) was from Jena Bioscience. All reactants and salts were of analytical grade (Merck). Chloroform was spectroscopic grade (Merck).

### Strains, plasmids and cell culture

All *E. coli* strains and plasmid used in this study are listed in Table 1. Cells were grown in Luria-Bertani (LB) medium at 30°C, 37°C or 42°C (as indicated) supplemented with the appropriate antibiotics (ampicillin 50 μg ml^-1^, chloramphenicol 15 μg ml^-1^ or tetracycline 10 μg ml^-1^) and gene expression inducers, IPTG (Isopropyl beta-D-1-thiogalactopyranoside) and sodium salicylate.

**Table 1.**
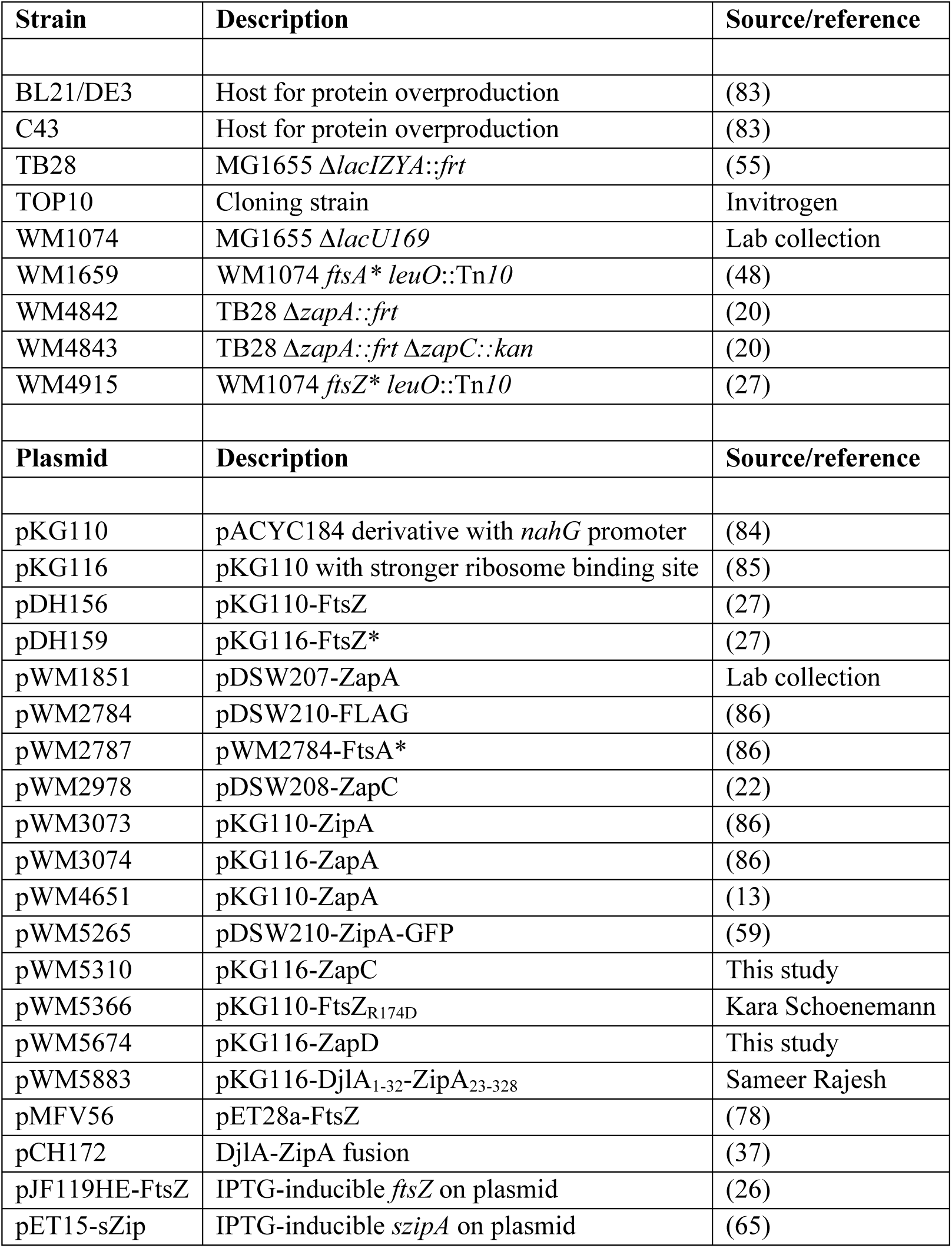
Strains and plasmids used in this study.

Overnight cell cultures were diluted 1:100 in the appropriate media and grown until OD_600_=0.2 followed by their back-dilution 1:4. After the second dilution, cells were cultured to OD_600_=0.2 and spotted on plates at 1x, 0.1x, 0.01x, 0.001x, and 0.0001x dilutions from right to left. For DIC microscopy they were further cultured in the presence of inducers, maintained in exponential phase, harvested 2h after induction and fixed with 1% formaldehyde.

### Plasmid constructions and DNA manipulation

Standard protocols for molecular cloning, transformation, and DNA analysis were used in this study (77). For cloning of DjlA_1-32_-ZipA_23-328_ in salicylate-inducible vector pKG116, we used the *djlA* forward primer (MK17: 5'-GGACTAGTATGCAGTATTGGGGAAAAATCATTGGC-3') and *zipA* reverse primer (MK18: 5-AAGGATCCTCAGGCGTTGGCGTCTTT-3), using pCH172 plasmid (37) kindly provided by Piet de Boer, as template. The cloning was confirmed by DNA sequencing.

### Protein purification and labeling

*E. coli* FtsZ was purified by the Ca^2+^-induced precipitation method (78). The soluble mutant of ZipA lacking the trans-membrane region (sZipA) was isolated as described (65). FtsZ and ZipA were labeled with Alexa probes (1:10 molar ratio). FtsZ was labeled under conditions that promote protein polymerization to ensure minimal interference of the dye with FtsZ assembly as described (79).

### Lipid monolayer assay

Lipid monolayers were prepared as described previously (13, 61). Briefly, 0.2 μg of *E. coli* polar lipid extract supplemented with 0.5-10% of NTA lipids when needed, were floated on Z-buffer (50 mM Tris HCl, pH 7.5, 300 mM KCl, 5 mM MgCl_2_) using a custom made teflon block (80) and placed in a humid chamber for 1h to evaporate the chloroform. Electron microscopy grids were then placed on the top of each well followed by sequential additions and incubations of 1 μM sZipA (1 h), 0.5-5 μM FtsZ (15 min) and 2 mM GTP or 0.5 mM GMPcPP (5 min). The grids were then removed followed by negative staining with uranyl acetate as described (27) and imaged with a JEOL 1230 electron microscope operated at 100 kV coupled with a TVIPS TemCam-F416 CMOS camera. FtsZ protofilament spacing was measured using the Plot Profile tool in ImageJ (81).

### Self-organization assays on supported lipid bilayers (SLBs)

Lipid bilayers were formed by fusion of small unilamellar vesicles (SUVs) mediated by CaCl_2_ (82). Lipids (polar extract phospholipids from *E. coli* or DOPC) with or without NTA at 0.5-1% w/w ratios, were prepared by drying a proper amount of the lipid stock solution under a nitrogen stream and kept under vacuum for at least 2 h to remove organic solvent traces. The dried lipid film was dissolved in SLB buffer (50 mM Tris-HCl, pH 7.5, 150 mM KCl) to a final 4 g/l concentration resulting in a solution containing multilamellar vesicles (MLVs). After 10 min sonication of MLVs, small unilamellar vesicles (SUVs) were obtained. One mg/mL suspension of SUVs was added to a hand-operated chamber (a plastic ring attached on a clean glass coverslip using UV-curable glue (Norland Optical Adhesive 63). SLBs were obtained by addition of 2 mM CaCl_2_ and incubated at 37°C for 20 min. Samples were washed with pre-warmed SLB buffer to remove non-fused vesicles.

Confocal images were collected with a Leica TCS SP5 AOBS inverted confocal microscope with a 63× (N.A. = 1,4–0,6/Oil HCX PL APO, Lbd.Bl.) immersion objective and Confocal multispectral Leica TCS SP8 system with a 3X STED (Stimulation Emission Depletion) module for super-resolution (Leica, Mannheim, Germany). TIRFM experiments were performed on a Leica DMi8 S widefield epifluorescence microscope. Images were acquired every 0.3 s with Hamamatsu Flash 4 sCMOS digital camera.

For self-organization assays, SLB buffer was replaced by Z-buffer prior to protein addition. The final volume of the assays was 100 μl. First, 0.5 μM of Alexa Fluor 647-labeled sZipA was added on top of the lipid bilayer with a given amount of NTA lipids. Once the fluorescent sZipA was visualized as attached to the lipid bilayer, 1 μM FtsZ-Alexa 488 was added followed by addition of 2 mM GTP (or 0.5 mM of GMPcPP) to induce FtsZ polymerization.

### Data availability

The authors declare that all data supporting the findings of this study are available within the article, Supplementary Information, or from the authors upon request.

## Acknowledgements

We thank Sameer Rajesh and Naga Babu Chinnam for strain constructions and preliminary experiments, Miguel Vicente and Ana Isabel Rico for the pJF119HE-FtsZ plasmid and Piet de Boer for the DjlA-ZipA fusion plasmid. We are grateful to members of the Margolin and Rivas laboratories for helpful discussions, and the Confocal Microscopy service at the Centro Nacional de Biotecnología, and the Electron Microscopy and the Confocal Microscopy services at the Centro de Investigaciones Biológicas, both at Consejo Superior de Investigaciones Científicas, for technical support. This work was supported by grant GM61074 from the National Institutes of Health to W.M. and by grant BFU2016-75471-C2-1-P from the Spanish Government to G.R.

## Supplemental Figure legends

**Figure S1. Immunoblots showing cellular levels of FtsZ and ZipA in various induction conditions and genotypes.** SDS-PAGE of cell extracts from (A) TB28 (WT), *ΔzapA* (Δ*A*) and *ΔzapAΔzapC* (*ΔAC*) backgrounds and (B) WM1074 (WT) and its derivatives containing chromosomal *ftsA\** and *ftsZ\** were blotted and probed with anti-FtsZ or anti-ZipA polyclonal antibodies (48). Relative band intensities for each of the 4 blots were analyzed and plotted in ImageJ, with the weakest band in each set normalized to 1.

**Figure S2. Excess ZipA cannot counteract the dominant negative effects of an under-bundled FtsZ.** WM1074 wild-type cells co-transformed with pKG110-FtsZ (p-FtsZ) or toxic, dominant negative FtsZ_R174D_ (p-FtsZ_R174D_) and pDSW210 (pEV) or pDSW210-ZipA-GFP (pZipA) were spotted on plates containing indicated concentrations of inducers to test if the toxicity of FtsZ_R174D_ could be antagonized by ZipA.

**Figure S3. sZipA does not form structures on lipid monolayers and FtsZ only forms residual sporadic filaments on monolayers supplemented with DGS-NTA.** sZipA (2 μM) was incubated on lipid monolayers containing 10% NTA lipids (A). 5 μM FtsZ was incubated on monolayers with 1% of NTA lipids, but not seeded with sZipA (B). Grids were negatively stained and visualized by electron microscopy. Arrow highlights a single FtsZ protofilament. Scale bar= 100 nm.

**Figure S4. Assembly of FtsZ on lipid monolayers containing sZipA and the effect of FtsZ concentration and of lipid composition.** (A) Circular structures of FtsZ single filaments on a DOPC monolayer containing 1% of DGS-NTA and 2 μM sZipA in the presence of GMPcPP. FtsZ concentration was 5 μΜ. (B) The equivalent experiment was performed on an *E. coli* polar lipid monolayer. Scale bars are shown.

**Figure S5. Kymographs of FtsZ swirls on SLBs carrying low-density sZipA.** Representative kymographs tracking the circumferential motion of individual FtsZ Alexa 488 swirls (as seen on videos) (A) with added GTP or (B) with added GMPcPP. Time and length scales are indicated, and black lines denote two representative paths of traveling fluorescence intensities. The slopes of the lines correspond to the velocity (x/t). Kymographs were obtained using an ImageJ kymograph plugin (Jens Rietdorf and Arne Seitz, EMBL, Heidelberg); 20 different circular structures were analyzed.

**Video S1: GTP-dependent assembly of FtsZ on supported *E. coli* lipid bilayers containing low-density sZipA.** Video of FtsZ ring-like filaments formation after GTP addition. FtsZ Alexa Fluor 488 and sZipA concentrations are the same as in Figure 6A.

**Video S2: GMPcPP-dependent assembly of FtsZ on supported *E. coli* lipid bilayers containing low-density sZipA.** Video of FtsZ ring-like filaments formation after GMPcPP addition. FtsZ Alexa Fluor 488 and sZipA concentrations are the same as in Figure 6B.

